## Supplementary Figures 1-10, Supplementary Tables 1-3 for "Genetic and karyotype divergence between parents affect clonality and sterility in hybrids"

**The PDF file includes:**

Figs. S1 to S10

Tables S1 to S3

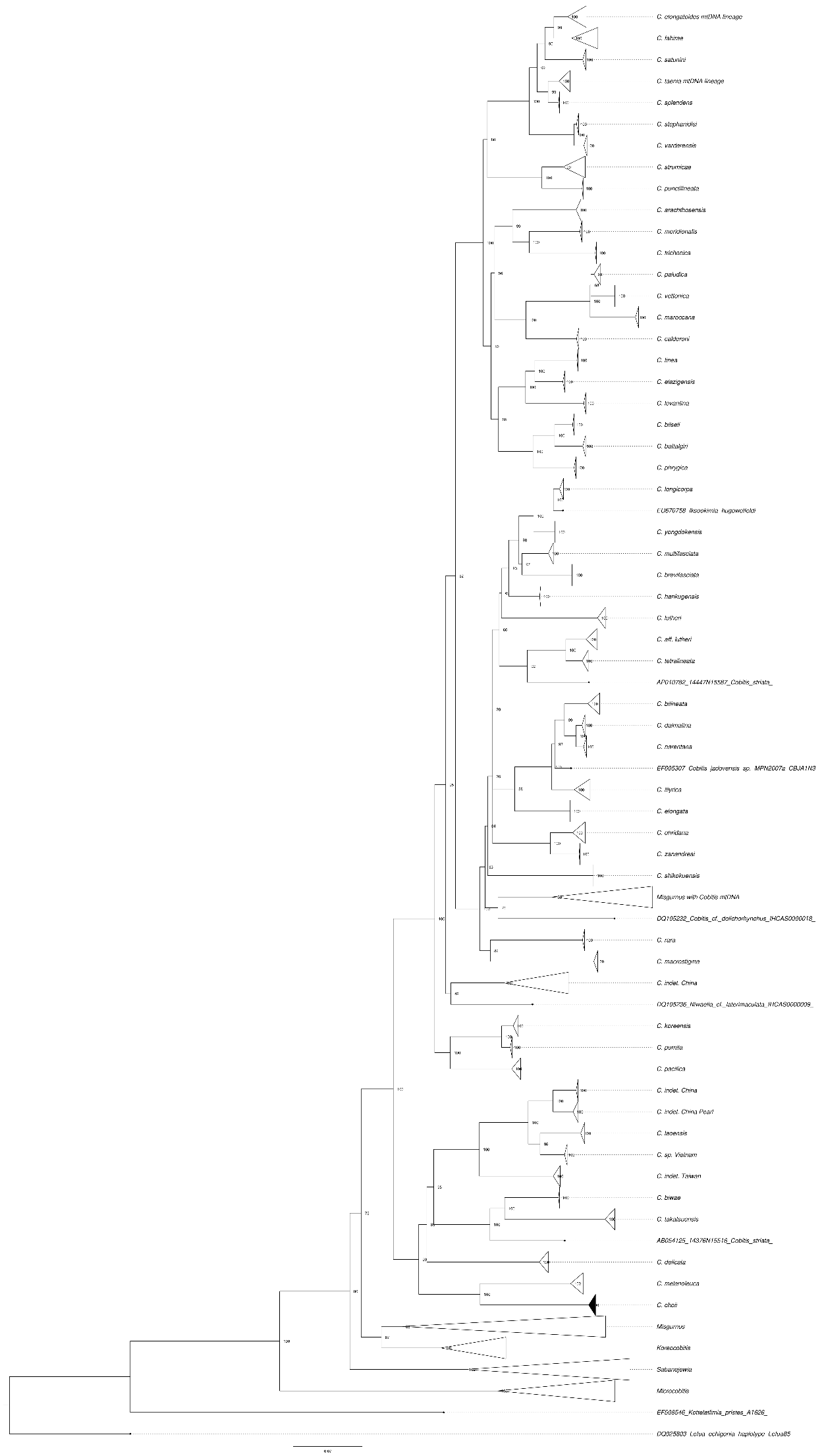

**C. hankugensis*

**C. lutheri*

**C. tetralineata*

***C. taenia***

***C. taurica***

***C. elongatoides***

***C. pontica***

***C. tanaitica***

**I. longicorpa*

***C. bilineata***

***C. ohridana***

**Misgurnus with Cobitis mtDNA*

*C. biwae*

*Misgurnus without mt introgression*

*Lefua echigonia*

***C. strumicae***

**Figures S1. Phylogenetic relationships of the Cobitidae family based on the cytochrome b dataset (modified version of Perdices and co-authors (2016)**^21^**.** The species used in the current work for artificial crosses were bold labeled. These species belongs to three major lineages which colonized independently European fresh waters (Perdices et al. 2016). *Cobitis* sensu stricto lineage (yellow) include all the known species of the *C. taenia* complex, which known to hybridize and produce asexual diploid and polyploid hybrids. Adriatic lineage (blue) is represented mainly by allopatric species, in this study is represented by *C. ohridana* and *C. bilineata*. Bicanestrini group (green) used in this study is represented by *C. strumicae*. Species used in the current studz are bolded. Additionally, species which were previously reported to interbreed are marked with asterisk; species which were artificially crossed in previous works are underlined, all species contain the following abbreviations: *C. biwae* = BW; *C. lutheri* = Lut; *C. tetralineata* = Tet; *M. anguillicaudatus* lineage with *Cobitis* mitochondrial introgression = MAmtC; *M.*

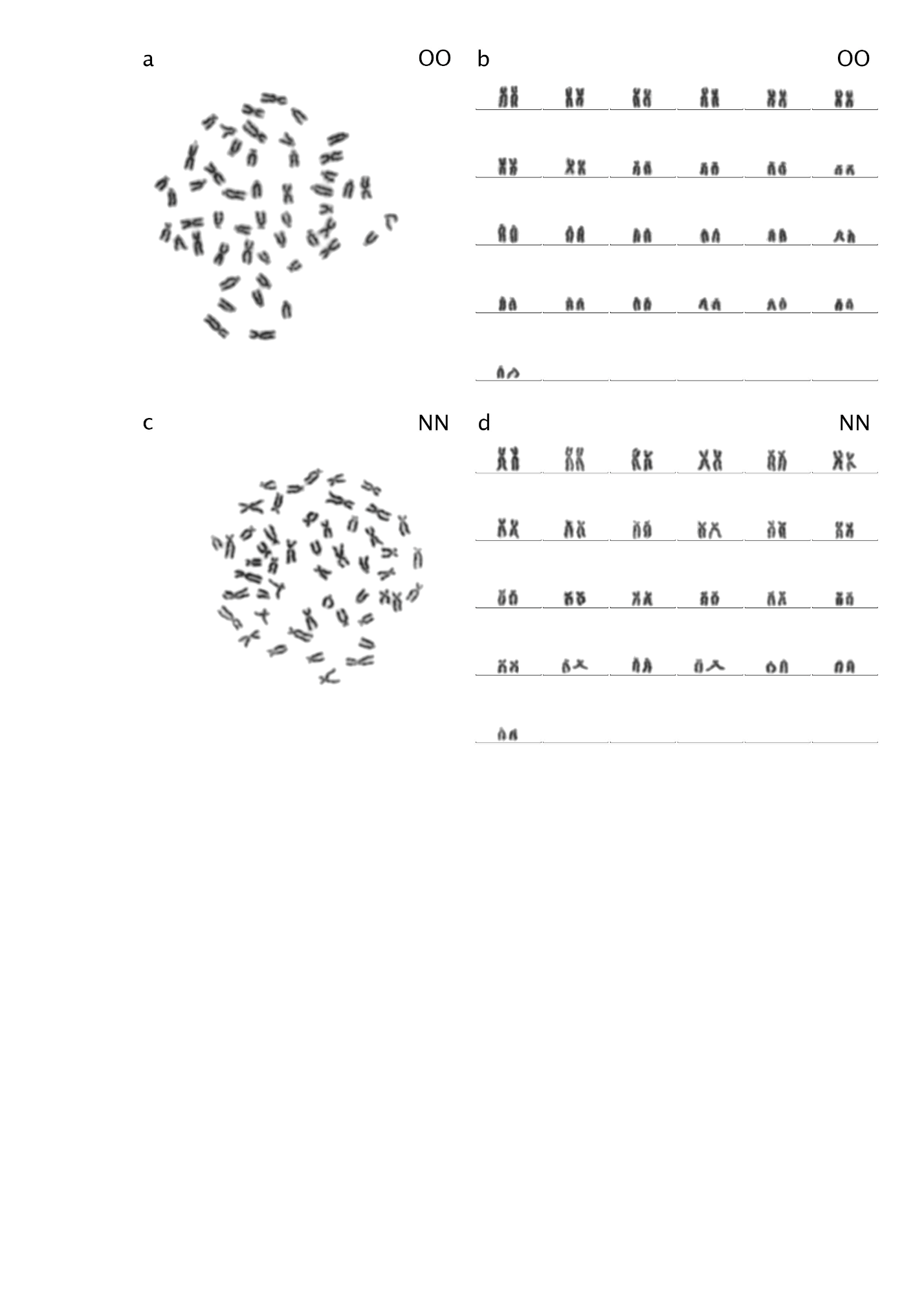

**Figure S2. Karyotypes and karyograms of *C. ohridana* (a,b) and *C. tanaitica* (c,d).** Both species have 2n = 50 chromosomes. *C. ohridana* has 20 meta-/submetacentric chromosomes and 30 subtelo-/ acrocentric chromosomes. *C. tanaitica* used in this study has 40 meta-/submetacentric chromosomes and 10 subtelo-/ acrocentric chromosomes. *C. ohridana* is abbreviated as OO; *C. tanaitica* is abbreviated as NN.

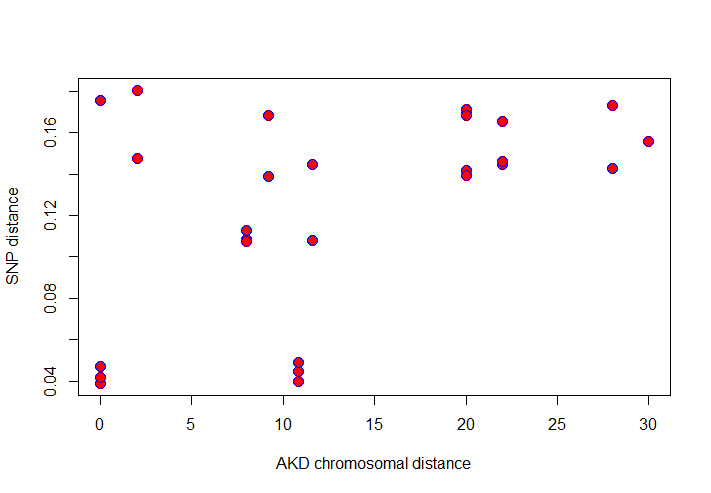

**Figure S3.** **Plot of correlation between Castigla’s AKD index (x-axis) and exome-wide genetic distance (SNP; y-axis).**

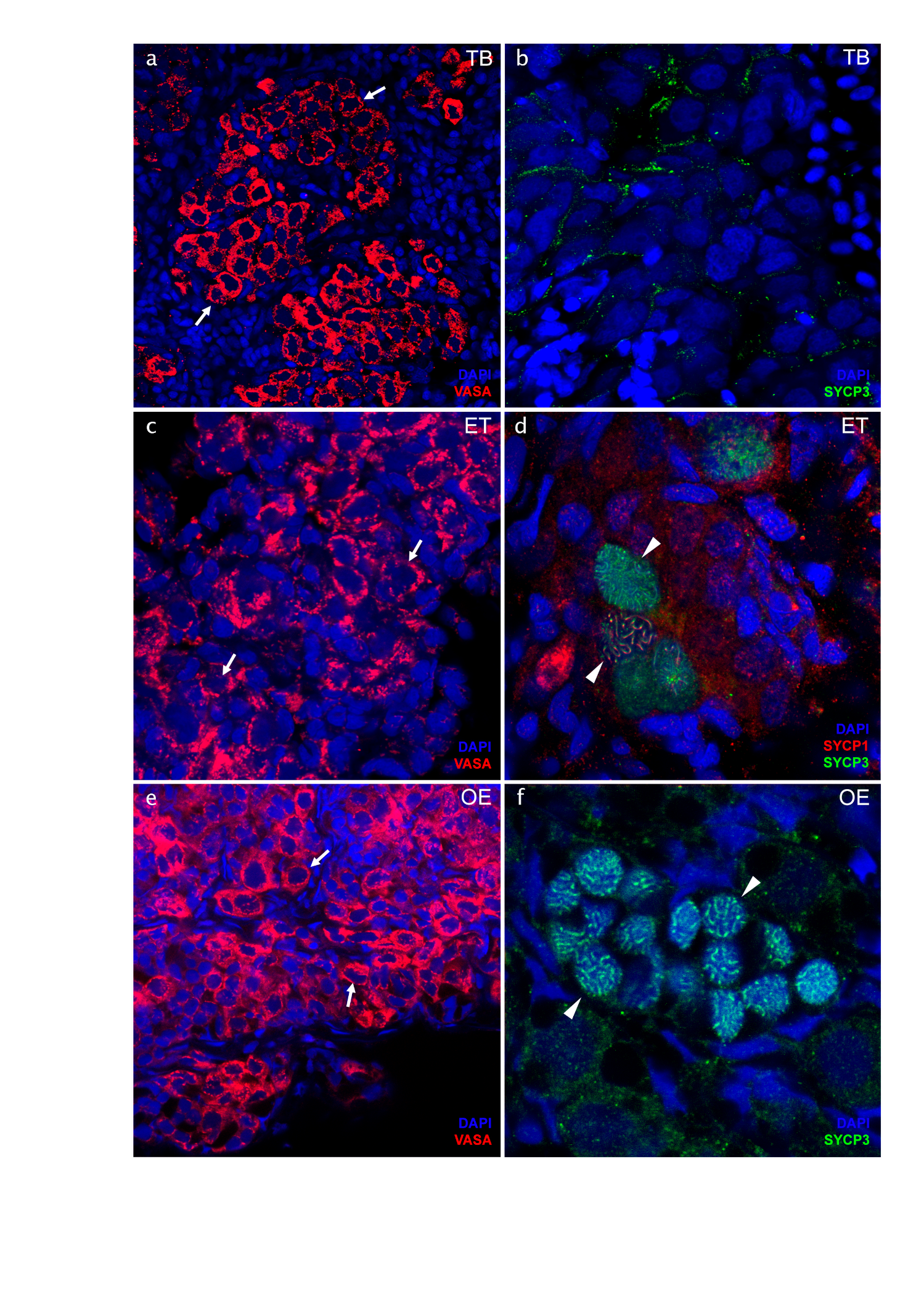

**Figure S4. Identification of germ cells and meiotic cells in the gonads of TB (a,b), ET (c,d) and OE (e,f) hybrids.** Germ cells identified using anti-vasa antibodies (indicated by arrows) were present in all hybrids (a,c,e); meiotic cells during pachytene (indicated by arrowheads) identified using anti SYCP3 and SYCP1 antibodies were not detected in TB hybrids while present in ET and OE hybrids (b,d,f). Scale bars = 10 µm.

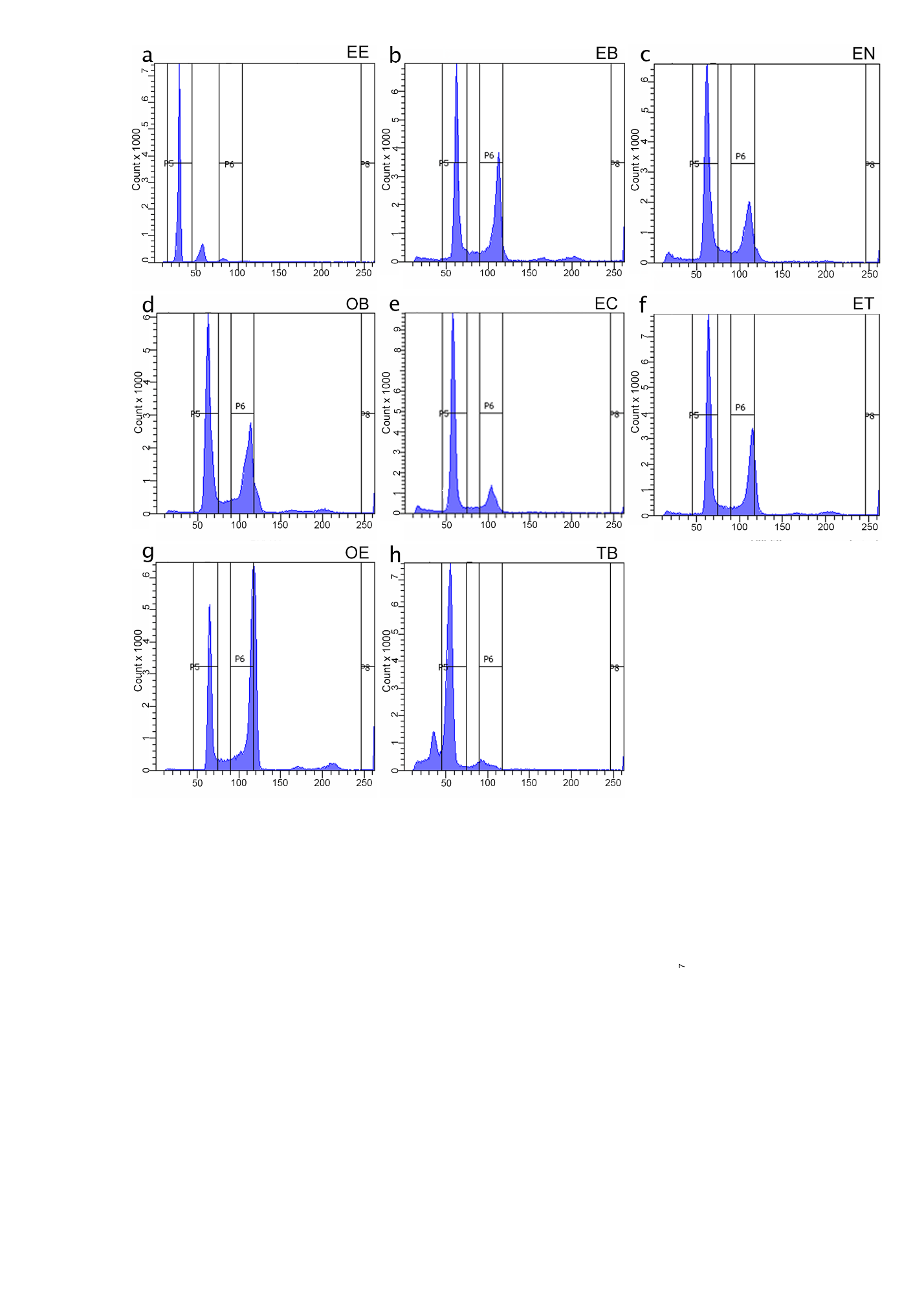

**Figure S5. Flow cytometry results of the analysis of cell suspension from testis of *C. elongatoides* (a), and diploid hybrid males (b-h).** (a) Sexual male has clear 1C cells corresponding to haploid sperm; diploid (2C) cell populations corresponding to spermatogonia; somatic cells and double-diploid (4C) cell populations corresponding to primary spermatocytes. (b-h) All hybrid males lack the peak of 1C cells but exhibit diploid (2C) and high amount of double-diploid (4C) cell populations accumulated possibly due to problems during chromosomal pairing during pachytene.

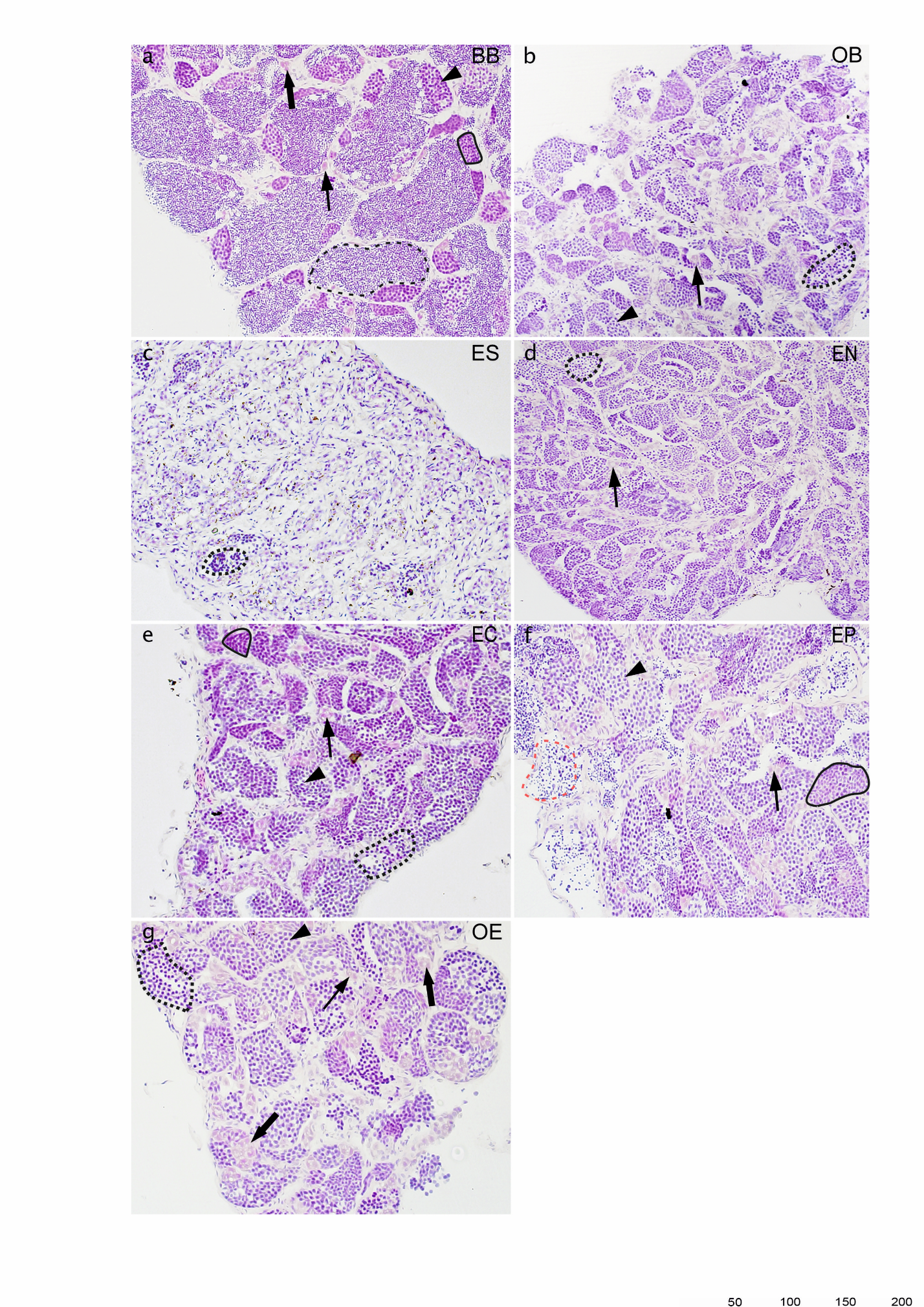

**Figure S6. Comparison of spermatogenesis between male of sexual diploid species *C. bilineata* (a) and diploid OB (b), ES (c), EN (d), EC (e), EP (f), OE (g) hybrids.** Semithin gonadal section of sexual *C. bilineata* male (a) shows spermatogonia type A (thin arrow), and spermatogonia type B (thick arrow), spermatocytes type I (arrowhead) and spermatocytes type II (black line), as well as spermatozoa (black dash line) in sexual males. Testes of hybrids (b-g) displays defective development with the presence of spermatogonia type A and type B, spermatocytes type I and spermatocytes type II (black line), but only a few spermatids (dotted black line) and/or aberrant spermatozoa (red dash line) germ cells of one cyst at different stages (black dotted line); (b) spermatogonia A with nucleus (N). Scale bar = 100 µm.

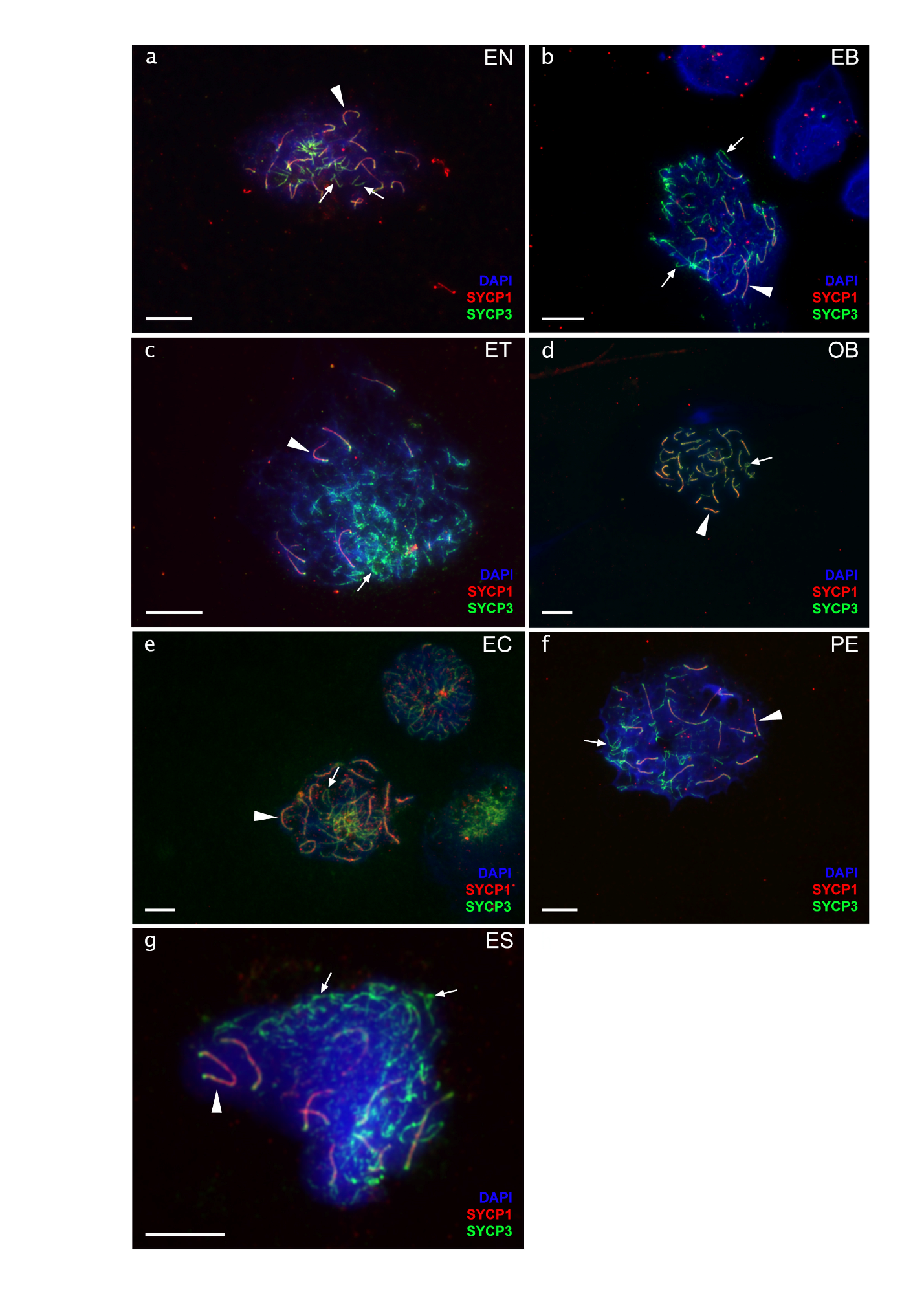

**Figure S7 Chromosomal spreads of pachytene spermatocytes of diploid EN (a), EB (b), ET (c), OB (d), EC (e), PE (f) and ES (g) males**. Synaptonemal complexes are visualized by immunolabeling with antibodies against SYCP3 protein (green) and SYCP1 (red) stained with DAPI (blue). Both SYCP3 and SYCP1 proteins are localized on bivalents (indicated by arrowhead) while only SYCP3 protein is localized on univalent (indicated by thin arrows). Pachytene spreads of diploid hybrid males have incomplete pairing with the number of bivalents and univalents. Scale bar = 10µm.

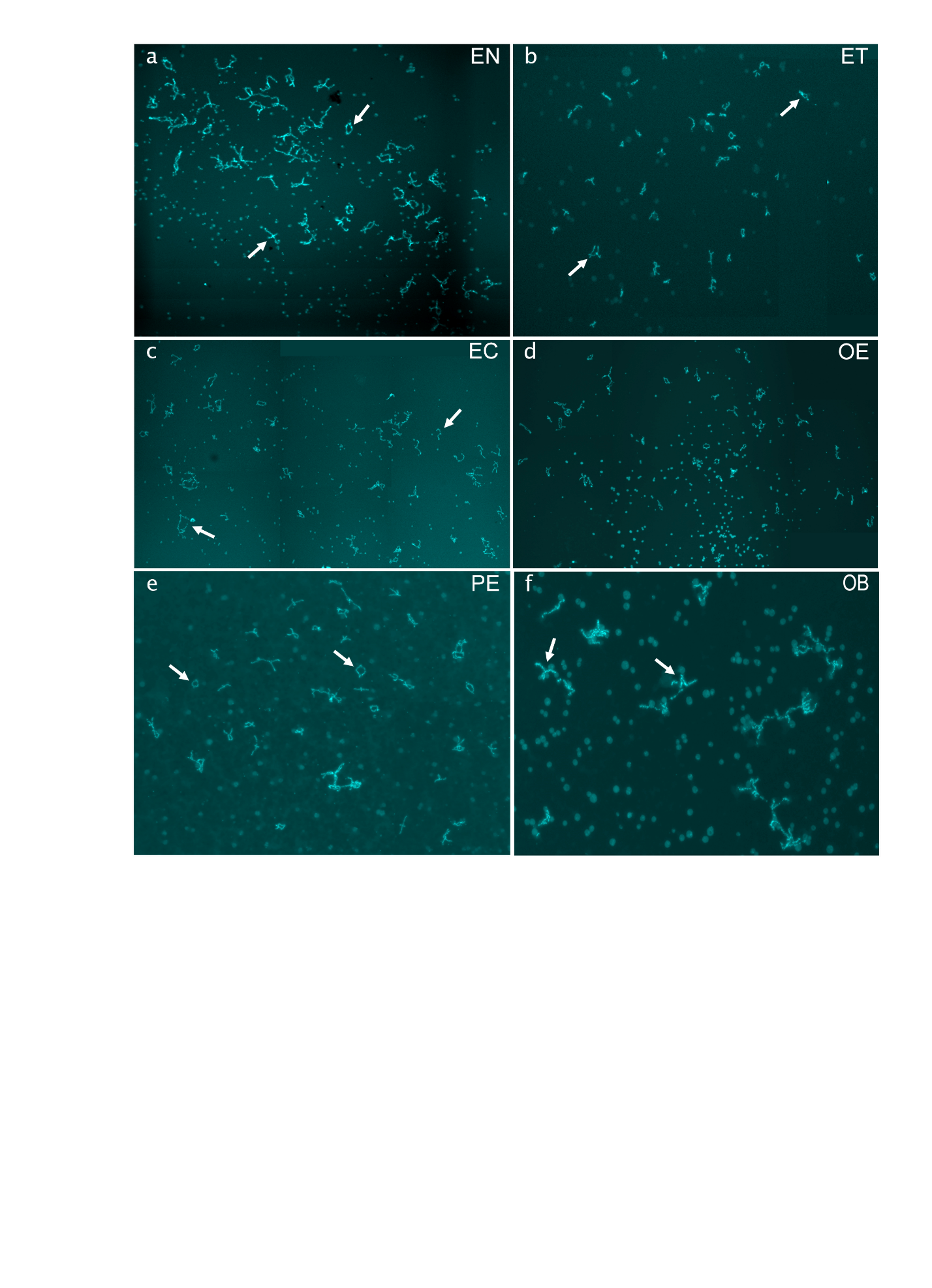

**Figure S8. Chromosomal spreads from diplotene oocytes of diploid EN (a), ET (b), EC (c), OE (d), PE (e) and OB (f) hybrid females.** Full diplotene chromosomal spread from the individual oocyte including 50 (a,c,d,e) and 49 (b) with no univalent or aberrant pairing, suggesting the pairing of homologous chromosome emerged after premeiotic endoreplication. Full diplotene chromosomal spread from OB hybrid is represented by 25 bivalents (f), suggesting the normal pairing of orthologous (O×B) chromosomes. Since the chromosomal spread from individual oocyte was large, several images were taken and merged into one. Arrows show examples of individual bivalents. Scale bar = 50 µm.

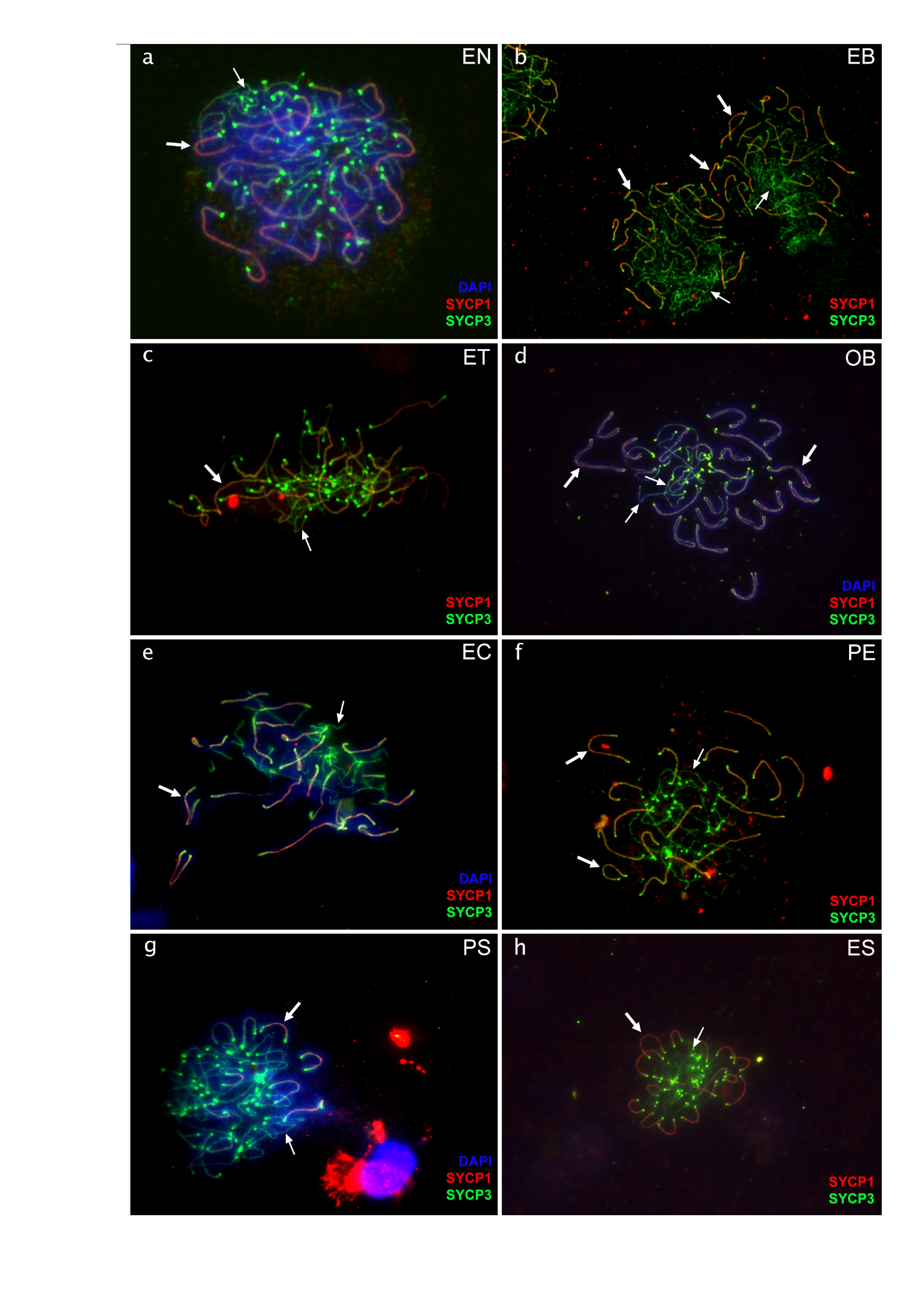

**Figure S9. Chromosomal spreads of pachytene oocytes with nonduplicated genomes of diploid EN (a), EB (b), ET (c), OB (d), EC (e), PE (f), PS (g) and ES (h) females**. Synaptonemal complexes are visualized by immunolabeling with antibodies against SYCP3 protein (green) and SYCP1 (red) stained with DAPI (blue). Both SYCP3 and SYCP1 proteins are localized on bivalents (indicated by arrows) while only SYCP3 protein is localized on univalent (indicated by arrowheads). Pachytene spreads from oocytes with unduplicated genome in diploid hybrid females exhibit incomplete pairing with the number of bivalents and univalent. Scale bar = 10µm.

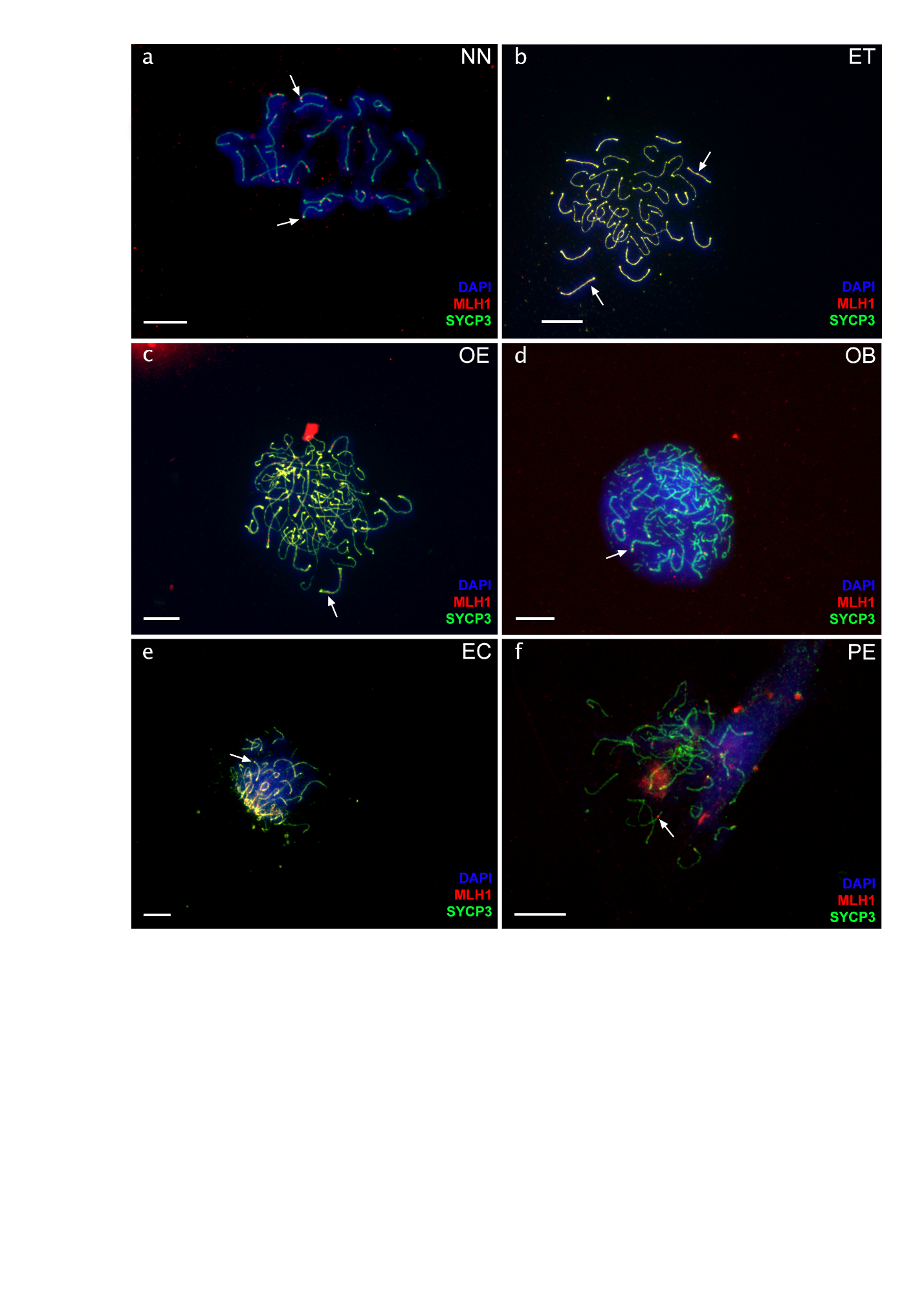

**Figure S10.** **Chromosomal spreads of pachytene oocytes of sexual species NN (a) and hybrids ET (b), OE (c), OB (d), EC (e) and PE (f) males**. Synaptonemal complexes are visualized by immunolabeling with antibodies against SYCP3 protein (green) and recombination loci were detected by antibodies against MLH1 protein (indicated by arrows, red). Chromatin stained with DAPI (blue). Recombination loci were observed only on bivalents, while univalents have no signal. Pachytene spreads from spermatocytes exhibit aberrant pairing. Scale bar = 10µm.

**Table S1.**

| Species ID | Nr. of chromosomes | Nr. of m/sm | Nr. of a/st | Ratio of a/st | Source of the karyotypes | RAG1 accession numbers |
| --- | --- | --- | --- | --- | --- | --- |
| *C. elongatoides* | 50 | 48 | 2 | 0.040 | *(69)* | KJ885726, KJ885727, KJ885728, KJ885730, KJ885731 |
| *C. taenia* | 48 | 28 | 20 | 0.417 | *(69)* | KJ885721, KJ885722, KJ885723, KJ885724, KJ885725 |
| *C. pontica* | 50 | 40 | 10 | 0.200 | Current study | CRUM3_1_pontica, CRUM3_1b_pontica, COB3_3_pontica |
| *C. tanaitica* | 50 | 40 | 10 | 0.200 | *(22)* | KJ885732, KJ885733 |
| *C. taurica* | 50 | 40 | 10 | 0.200 | *(70)* | COBVB2_3_taurica,COBVB2_3b_taurica, COBVB13_3_taurica, COBVB13_2_taurica |
| *C. strumicae* | 50 | 20 | 30 | 0.600 | *(71)* | KP161179, KP161180 |
| *C. lutheri* | 50 | 16 | 34 | 0.680 | *(72)* | JN858843, JN858844, KP161170, KP161171, KP161172 |
| *C. tetralineata* | 50 | 16 | 34 | 0.680 | *(72)* | KP161183, KP161184, KP161185 |
| *M. anuguillicaudatus*_clade B | 50 | 14 | 36 | 0.720 | *(73)* | AB698049, AB698050, AB698057, AB698058, AB698059, AB698060, AB698061, AB698062, AB698063, AB698064 |
| *C. hankugensis* | 48 | 18 | 30 | 0.625 | *(74)* | MH464460, MH464461, MH464462, MH464463, MH464464, MH464465, MH464466, MH464467, MH464468, MH464469, MH464470, MH464471, MH464472, MH464473, MH464474, MH464475 |
| *C. bilineata* | 50 | 18 | 32 | 0.640 | *(75)* | EF672416, KP161139 |
| *C. ohridana* | 50 | 20 | 30 | 0.600 | Current study | EF672431, KP161161, EF672432, EF672433 |
| *I. longicorpa* | 50 | 16 | 34 | 0.680 | *(74)* | MH464520, MH464521, MH464522, MH464523, MH464524, MH464525, MH464526, MH464527, MH464528, MH464529, MH464530, MH464531, MH464532, MH464533 |
| *C. biwae* | 48 | 38 | 10 | 0.208 | *(76)* | AB531331, AB531336, AB531337 |
| *M. anguillicaudatus*_clade A | 50 | 14 | 36 | 0.720 | *(73)* | AB698051, AB698052, AB698053, AB698054, AB698055, AB698056 |
| *L. echigonia* | 50 | 12 | 38 | 0.760 | *(77)* | DQ025803 |

**This study uses a summary of karyotypes and RAG1 sequence of loaches (Cobitidae).** The references are indicated for species karyotypes that were previously described. The Karyotype of *C. ohridana* is published for the first time in this work.

**Table S2.**

| Crosstype | Nr. of families | Family ID | Nr. of obtained babies  per family | Nr. of females and  males per family | | Undetermined sex | Confocal analysis of germ cells using vasa | Analysed specimens for pachytene | Analysed specimens for diplotene | Flow- cytometry | Histology |
| --- | --- | --- | --- | --- | --- | --- | --- | --- | --- | --- | --- |
| O x E | 2 | Cobex_25 | 6 | ♀ | 5 | 0 | 1 | 3 | 0 | 0 | 0 |
|  |  |  |  | ♂ | 1 |  | 0 | 0 | 0 | 0 | 1 |
|  |  | Cobex_30 | 30 | ♀ | 15 | 2 | 2 | 4 | 2 | 0 | 0 |
|  |  |  |  | ♂ | 13 |  | 2 | 3 | 0 | 2 | 1 |
| E x O | 1 | Alex_45 | 59 | ♀ | 23 | 14 | 2 | 3 | 2 | 0 | 0 |
|  |  |  |  | ♂ | 22 |  | 2 | 2 | 0 | 2 | 0 |
| O x B | 2 | Cobex_24 | 25 | ♀ | 10 | 8 | 1 | 3 | 2 | 0 | 0 |
|  |  |  |  | ♂ | 7 |  | 1 | 1 | 0 | 1 | 2 |
|  |  | Cobex_29 | 19 | ♀ | 7 | 6 | 1 | 1 | 1 | 0 | 0 |
|  |  |  |  | ♂ | 6 |  | 1 | 2 | 0 | 1 | 2 |
| P x S | 1 | 2017PS | 17 | ♀ | 11 | 0 | 0 | 4 | 2 | 0 | 0 |
|  |  |  |  | ♂ | 6 |  | 2 | 4 | 0 | 0 | 0 |
| P x E | 1 | 2017EP | 20 | ♀ | 10 | 1 | 0 | 5 | 2 | 0 | 0 |
|  |  |  |  | ♂ | 9 |  | 0 | 2 | 0 | 0 | 2 |
| E x S | 1 | 2017ES | 28 | ♀ | 7 | 0 | 0 | 6 | 4 | 0 | 0 |
|  |  |  |  | ♂ | 21 |  | 0 | 2 | 0 | 0 | 3 |
| E x B | 2 | Cobex11 | 10 | ♀ | 7 | 1 | 0 | 1 | 0 | 0 | 0 |
|  |  |  |  | ♂ | 2 |  | 0 | 1 | 0 | 1 | 0 |
|  |  | 2020EB | 3 | ♀ | 1 | 0 | 0 | 0 | 0 | 0 | 0 |
|  |  |  |  | ♂ | 2 |  | 0 | 1 | 0 | 1 | 0 |
| E x T | 3 | Cobex_12 | 5 | ♀ | 3 | 0 | 0 | 1 | 1 | 0 | 0 |
|  |  |  |  | ♂ | 2 |  | 1 | 1 | 0 | 0 | 0 |
|  |  | 2018ET | 35 | ♀ | 11 | 16 | 1 | 1 | 2 | 0 | 0 |
|  |  |  |  | ♂ | 8 |  | 0 | 1 | 0 | 1 | 0 |
|  |  | Alex_46 | 33 | ♀ | 8 | 15 | 1 | 1 | 1 | 0 | 0 |
|  |  |  |  | ♂ | 10 |  | 1 | 1 | 0 | 1 | 0 |
| T x E | 1 | Alex_65 | 15 | ♀ | 6 | 2 | 2 | 2 | 2 | 0 | 0 |
|  |  |  |  | ♂ | 7 |  | 2 | 1 | 0 | 3 | 0 |
| T x B | 1 | 2020TB | 16 | ♀ | 4 | 1 | 0 | 1 | 0 | 0 | 0 |
|  |  |  |  | ♂ | 11 |  | 3 | 3 | 0 | 2 | 0 |
| E x C | 1 | Cobex_37 | 30 | ♀ | 11 | 11 | 0 | 3 | 3 | 0 | 0 |
|  |  |  |  | ♂ | 8 |  | 0 | 3 | 0 | 3 | 2 |
| E x N | 2 | Alex_47 | 41 | ♀ | 20 | 3 | 1 | 4 | 1 | 0 | 0 |
|  |  |  |  | ♂ | 18 |  | 1 | 2 | 0 | 3 | 1 |
|  |  | 2020AF_13 | 23 | ♀ | 7 | 8 | 1 | 1 | 1 | 0 | 0 |
|  |  |  |  | ♂ | 8 |  | 1 | 2 | 0 | 0 | 1 |
| N x P | 1 | 2021NP | 34 | ♀ | 18 | 0 | 0 | 2 | 0 | 0 | 0 |
|  |  |  |  | ♂ | 16 |  | 0 | 1 | 0 | 0 | 0 |

**The numbers of analyzed male and female *Cobitis* samples from each F1 obtained family.** Most specimens were used for the pachytene analysis in order to increase the chance to observe duplicated cells in females, as their incidence per slide is very low in clonal hybrids. Also, we used as many individuals as possible for this approach to have a statistically significant number of SC per individual.

**Table S3.** Dataset with every SC reported from all analyzed specimens. **All this information was used in order to generate Figure 5.**
